# Evaluation of Convolutionary Neural Networks Modeling of DNA Sequences using Ordinal versus one-hot Encoding Method

**DOI:** 10.1101/186965

**Authors:** Allen Chieng Hoon Choong, Nung Kion Lee

## Abstract

Convolutionary neural network (CNN) is a popular choice for supervised DNA motif prediction due to its excellent performances. To employ CNN, the input DNA sequences are required to be encoded as numerical values and represented as either vectors or multi-dimensional matrices. This paper evaluates a simple and more compact ordinal encoding method versus the popular one-hot encoding for DNA sequences. We compare the performances of both encoding methods using three sets of datasets enriched with DNA motifs. We found that the ordinal encoding performs comparable to the one-hot method but with significant reduction in training time. In addition, the one-hot encoding performances are rather consistent across various datasets but would require suitable CNN configuration to perform well. The ordinal encoding with matrix representation performs best in some of the evaluated datasets. This study implies that the performances of CNN for DNA motif discovery depends on the suitable design of the sequence encoding and representation. The good performances of the ordinal encoding method demonstrates that there are still rooms for improvement for the one-hot encoding method.

## I. Introduction

CNN (Convolutional Neural Network) [1], [2] is currently one of the most widely used deep learning methods in machine learning due to its powerful modelling capability on complex and large-scale datasets. Recently, CNN has been widely used for learning DNA sequence datasets related to regulatory regions and other functional landmarks [3]–[6]. The advantage of CNN is its learning can be performed without the need of engineered features. The intrinsic features in the raw dataset are learned through the many layers structure which represents the different abstraction of features. The layers in a CNN consist of convolutionary and pooling layers. A convolutionary layer consists of multiple maps of neurons which are called filters. A filter convolves the inputs from the previous layer to produce a reduced sample. It only connected to a patch of the previous layer, which is named as “receptive field”. Moreover, all neurons in the filters detect the same features of the previous layer but at different map locations. Different filters might detect different types of features [7]. In a DNA dataset, the features might represent different motifs enriched in the input DNA sequences. In addition, [7] stated that the exact locations and frequency of a feature are unimportant to the learning purpose because the final output of the deep learning is recognition of the input data. On the other hand, the pooling layer summarizes the adjacent neurons by computing their activity. As a result, the model parameters are greatly reduced. After the last pooling layer, it has a fully connected multi-layers perceptron neural networks.

CNN is designed to effectively models multi-dimensional input data. Thus, it is powerful in solving problems related to computer vision and image recognition [2] where the data consists of images. To employ the CNN on DNA datasets, existing works typically encode the nucleotides in DNA sequences by using the one-hot method [3], [5], [6]. That is, each nucleotide is encoded with a binary vector of four bits with one of them is hot (i.e. 1) while others are 0. For instance *A* = (1,0,0,0), *G* = (0,1,0,0), *C* = (0,0,1,0), and *T* = (0,0,0,1). This sequence encoding method draws similarity to the Position Frequency Matrix [8]. In which, the values in a vector are considered as the probability of finding the four bases at a certain position in a DNA sequence. Once encoded, an input DNA sequence of length *l* is represented as 4 × *l* matrix. Or in another word, a “2D image” with one channel.

Methods for converting biological sequences into numerical values have been existed in numerious past studies [9]. Those encoding methods can be categorized into direct and indirect encoding [9]. Direct methods represent each nucleotides/amino acids with a numerical value or vector of numerical values. They preserved the original order the bases appeared in a biological sequence after the encoding. While the indirect methods engineered a fixed number of features (numerical values) from the biological sequences. The features can be based on frequency counts of various k-mers (short sequence segments of length *k* bp), biological, or biochemical properties.

In recent years, CNN and other deep learning techniques are gaining popularity in supervised DNA motif prediction because of their excellent performances. For example, in DeepBind, it achieved an average of 0.85 AUC for evaluation of 137 transcription factors, while MEME-Chip achieved only 0.82. Some of the recent works of using deep learning neural networks for DNA motif discovery are DeepBind [3], DeepSEA [6], Basset [5], TFImpute [10], and FIDDLE [11]. All of those works are using one-hot encoding to encode the input DNA sequences.

In this study, we propose a simple method to transform DNA sequences into matrix representation. The matrices are fed as input to CNN for classification model construction. We evaluate our propose encoding method to the most popular one-hot encoding method using the three sets of sequence datasets. This paper is organized as follows. The next section presents our method of this study including datasets used, evaluation metric, and CNN structure. The Results section presents our evaluation results with discussion. The last section discusses the main findings and some issues for future studies.

## II. Method

### A. Ordinal encoding

In CNN, the input examples are assumed to be represented as vectors or matrices of numerical values. In this study, we evaluate the utility of using ordinal encoding method to encode each nucleotide letter and thus a DNA sequence as a vector of numerical values. One of the problems with the one-hot encoding is the curse-of-dimensionality [12], in which for each input sequence of length *l*, the number of input values would be 4 × *l*. It causes long training time and only short sequence is computationally feasible for large-scale dataset. For examples, in DeepBind the sequence length is limited to 14101bp long, while Basset uses 600bp input sequences. While the one-hot encoding has been claimed to represent the PFM [3] and has a clear meaning of interpretation, for machine learning, that is unnecessary useful since our aim is to learn the features associated with different class labels. In our method, to reduce the dimensions of input sequence the nucleotides are encoded with numerical values. That is, A is represented by 0.25, C by 0.50, G by 0.75, and T by 1.00, respectively. For the unknown nucleotide N, its value is 0.00. It is difficulty to justify how those numbers are decided rather than with heuristic. But our preliminary evaluation indicated that those are good choices (results not shown).

The advantages of ordinal encoding is it requires less computer memory resource and computations. Moreover, a one-dimensional vector can be arbitrarily reshaped to a two-dimensional matrix. Suppose we have a row vector *v* of size 1 × *l*, where *l* is an even integer, that represents an input sequence. To transform a vector to a matrix of size *m* × *n*, we simply reshape the vector *v* to *m* × *n* matrix. If *l* is not multiple of *n*, zero values are padded at the last row.

### B. Datasets

We have prepared three sets of datasets to compare the ability of ordinal and one-hot encoding method to learn the motif features enriched in the DNA sequences. The datasets from Pazar database [13] are chosen for our comparative evaluation. Pazar is an open-access and open-source database that stores the transcription factor and regulatory sequence annotation independently [13]. The datasets are experimentally validated TFBSs. A set of mouse TF datasets were gathered from Pazar. The details of the datasets are shown in Table I. Another set of datasets collected from Pazar is the transcription factors of human. The information of the datasets are shown in Table II

**Table I.** Information of mouse datasets collected from Pazar

| Dataset | GEO | Number of sequences | Average length of sequences | File size |
| --- | --- | --- | --- | --- |
| BCL6 | GSE16723 | 1906 | 200.00 | 430.6k |
| CBP | GSE29362 | 6372 | 89.96 | 738.5k |
| CDX2 | GSE14586 | 59097 | 500.00 | 31.1M |
| CRX | GSE20012 | 9661 | 182.47 | 2.0M |
| ERG | GSE22178 | 36167 | 398.00 | 15.3M |
| ETS1 | GSE29362 | 2359 | 127.22 | 361.5k |
| EZH2 | GSE18776 | 4502 | 1449.55 | 6.6M |
| FLI1 | GSE22178 | 19601 | 398.00 | 8.3M |
| GATA2 | GSE22178 | 9234 | 398.00 | 3.9M |
| GFI1B | GSE22178 | 8853 | 398.00 | 3.8M |

**Table II.**
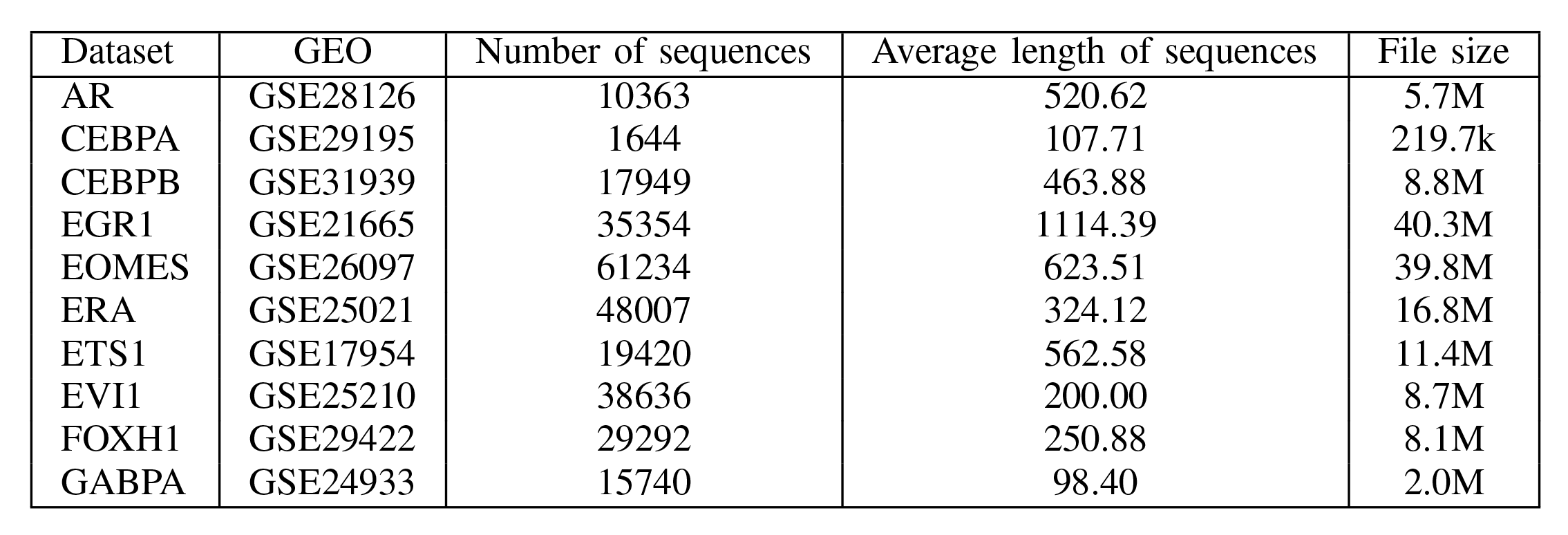
Information of human datasets collected from Pazar

Besides the datasets from Pazar, seven ChIP-seq datasets used in ENSPART [14] are also been selected (NRSF,FOXA1, CREB, FOXA2, OCT4, CTCF, STAT1).

To avoid classes imbalance problem, same number of sequences from each dataset is sampled. Furthermore, the sequences are truncated by removing bases that are beyond 900 bp. In our preliminary study (results not shown), using longer sequence lengths only gives marginal improvement on the accuracy rates but at the price of higher computational cost. Sequences that are shorter than 900 bp are padded with the vector [0.25,0.25,0.25,0.25] for the one-hot encoding, while 0.00s are added for the ordinal encoding.

Table III shows the number of samples from each dataset, number of sequences as training sets, validation sets, and testing sets.

**Table III.**
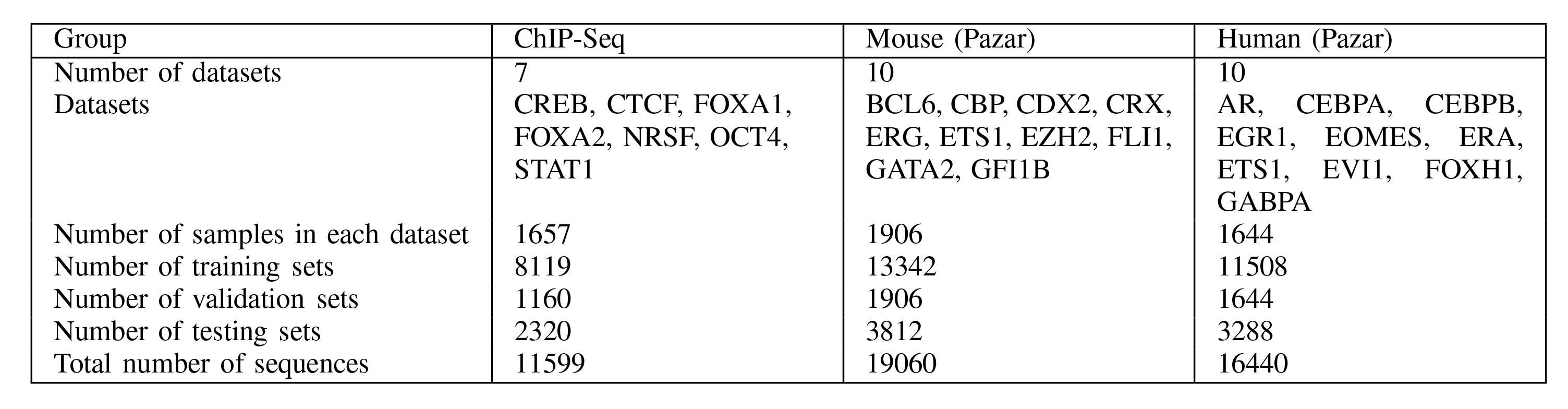
Number of samples for training, validation, and testing for each TF dataset.

### C. CNN Architecture

The evaluation study compares three (3) sequence representation using the two encoding methods:(a) one-hot with matrix representation; (b) ordinal encoding with square matrix (Square) representation; and (c) ordinal with 1-dimensional vector representation (1D). The CNN architectures employed in our simulation are are shown in Table IV. The CNN labeled “3Layer” is using three layers CNN with ordinal encoding and square matrix representation.

**Table IV.** CNN’s layers setup used in the simulations.

|  | Onehot | Square | 1D | 3layer |
| --- | --- | --- | --- | --- |
| Input layer | $900 \times 4$ | $30 \times 30$ | $900 \times 1$ | $30 \times 30$ |
| 1st conv layer | $12 \times 4$ | $6 \times 6$ | $36 \times 1$ | $6 \times 6$ |
| 1st activation function | ReLU | ReLU | ReLU | ReLU |
| 1st max pooling | $2 \times 2$ | $2 \times 2$ | $2 \times 2$ | $2 \times 2$ |
| 2nd conv layer | $8 \times 4$ | $6 \times 6$ | $36 \times 1$ | $6 \times 6$ |
| 2nd activation function | ReLU | ReLU | ReLU | ReLU |
| 2nd max pooling | $2 \times 2$ | $2 \times 2$ | $2 \times 2$ | $2 \times 2$ |
| 3rd conv layer | | | | $4 \times 4$ |
| 3rd activation function |  |  |  | ReLU |
| 3rd max pooling | | | | $2 \times 2$ |
| Fully connected layer | 1024 | 1024 | 1024 | 1024 |
| Dropout | 0.5 | 0.5 | 0.5 | 0.5 |
| Output layer | one-hot | one-hot | one-hot | one-hot |

“Square” representation uses a 30 × 30 input layer and “1D” uses 900 × 1 as a one-dimensional input layer. In addition, we also uses a three-layers CNN with the 30 × 30 square matrix with ordinal encoding. All the CNNs employed the dropout rate of 0.5. Dropout is applied before the output layer to reduce overfitting of training [15]. All the layers use ReLU activation function as recommended by [16].

The TensorFlow [17] is chosen as implementation of the CNN. The softmax activation function is used at the output layer. ADAM optimizer with the learning rate 10^−4^ is chosen as the adaptive learning method. The mini-batch of 200 is used in each epoch. Maximum epoch is set at 20000.

### D. Evaluation Metric

The Area under curves (AUC) [18] is used as the performance metric for the evaluation. We compare the one-hot and ordinal encoding methods using the tensor-flow implementation of the CNN. In addition, we also compare with Basset and gkm-SVM. A five-fold cross-validation is used for all the datasets. Basset and gkm-SVM do out AUC values in their output. While for the CNN implemented by the TensorFlow, the AUC values are computed by our own implementation.

## III. Results

Table V shows the comparison of average AUC rates from 5-fold cross-validation using the ChIP-seq datasets. It is observed that the square encoding method performed better than one-hot encoding for all the datasets. That is a surprising result since the one-hot has been the state-of-the-art encoding method for DNA sequences in most of the recent CNN works. It is also noted that the CNN with the square matrix performed better than gkm-SVM for 4 out of the 7 datasets. Other than that, Basset performed only better than CNN with square encoding in 2 of the 7 datasets. The results indicate that the two layers CNN might not be the most optimal architecture for the one-hot encoding use. However, the 3 convolutionary-pooling layers used by Basset is not better than the square encoding with 2 layers. Another observation is that CNN-based methods performed better that sVM in most of the evaluated datasets. This shows that CNN is more powerful in feature learning since it is able to learn the different abstraction of the features through its convolutionary-pooling layers. gkm-SVM employed k-mer feature for the modeling. To give a clearer comparison, Fig. 1 shows the graph of the average AUC values for all the datasets.

**Table V.**
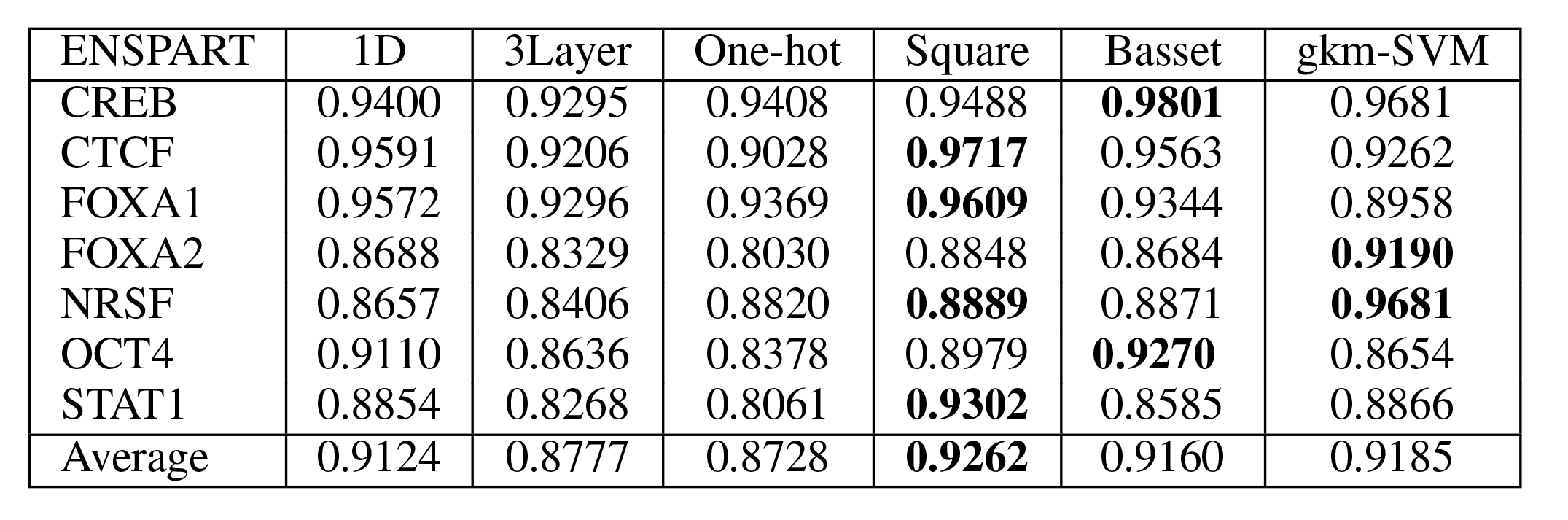
Comparison of the aucs of 1d, 3layer, onehot, and square of the average of 5-fold aucs with basset and gkm-SVM on chip-seq datasets

**Fig. 1.**
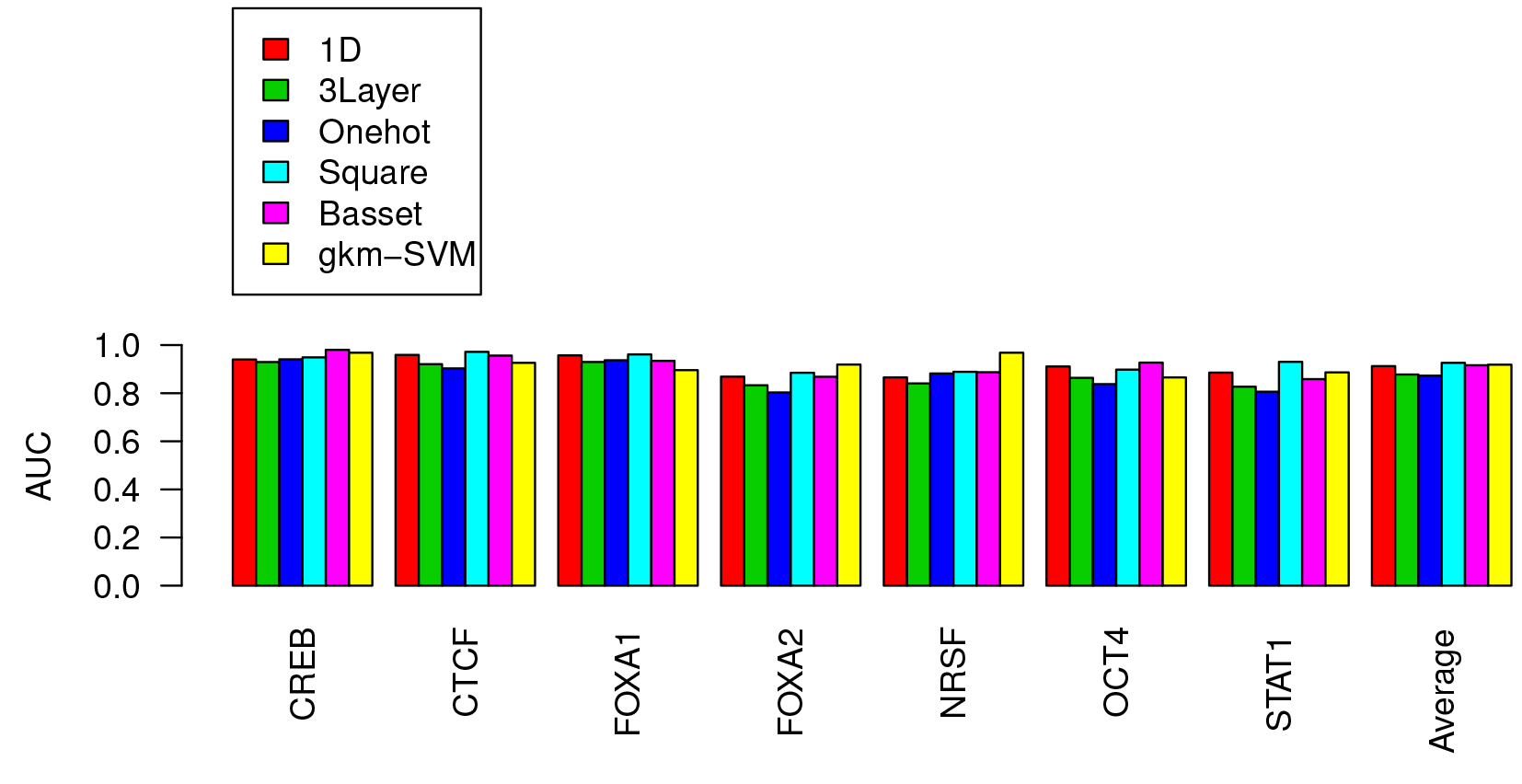
Comparisons of the CNN average AUCs using the 1D, one-hot, and square encoding. The AUC values are obtained from average of 5-fold crossvalidation using the ChIP-seq datasets.

Table III shows the comparisons of AUC values of CNN, Basset, and gkm-SVM using ordinal or one-hot encoding for the mouse datasets. It is noted that Basset performed better than other methods (gkm-SVM, CNN with ordinal encoding) in 9 out of 10 of the datasets. It obtained an average AUC value of 0.949 for ten of the datasets. However, it is noted that the CNN with square encoding produced AUC values which are quite close to the Basset’s performance (average AUC value on 10 datatasets is 0.92) and significantly better than gkm-SVM for 6 out of the 10 datasets. We also noted that the architecture used by the CNN might not be the most optimal because the one-hot encoding results should be quite close to the Basset since they are using the same encoding and representation. This highlights the difficulty of benchmarking CNN using different representations since their performances are also affected by the architecture and parameter values choices. It is also surprising to see that the AUC values of CNN using the 1D sequence encoding are very close to Basset. Fig. 2 illustrates the average AUC values for the mouse datasets.

**Fig. 2.**
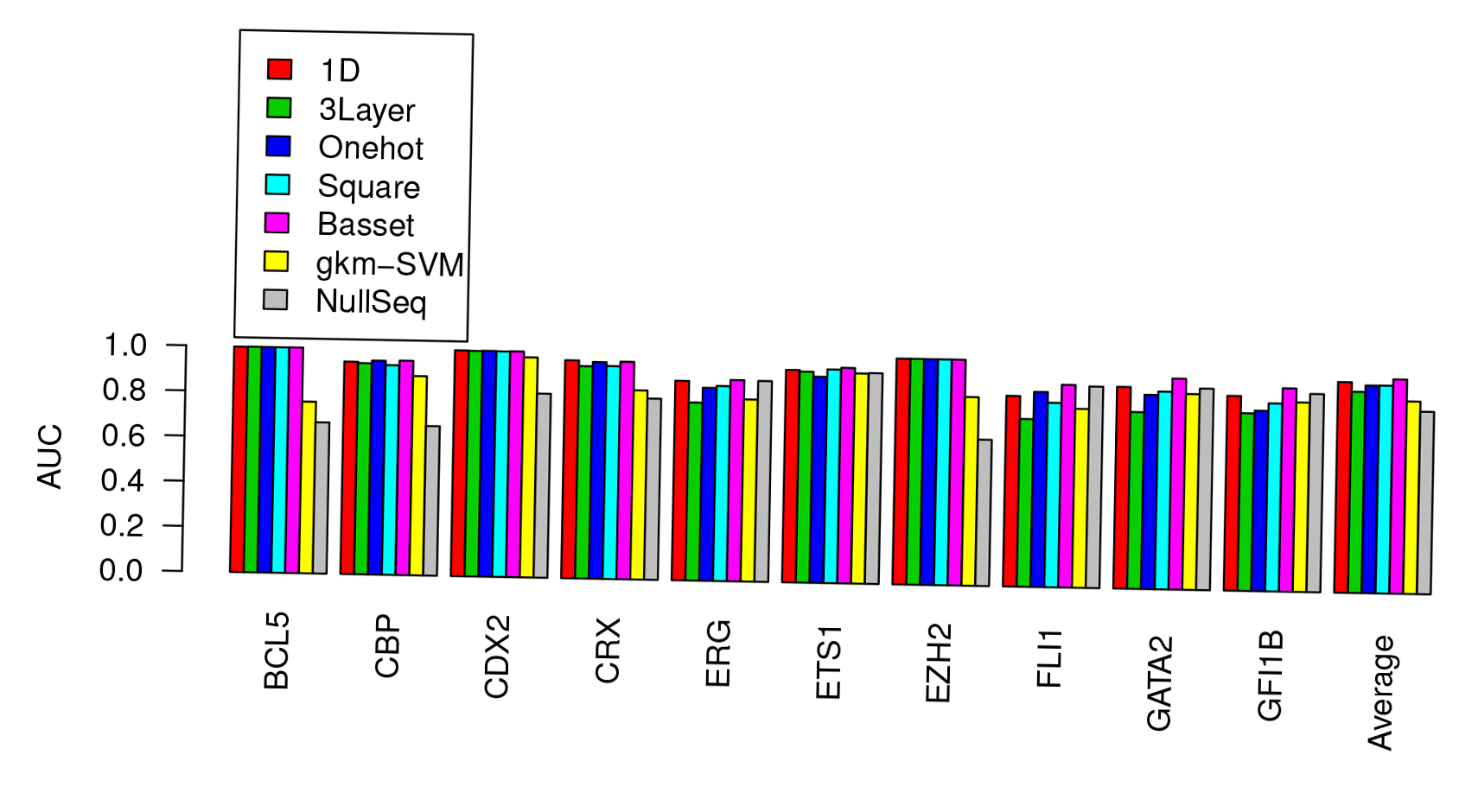
Comparison of the CNN average AUCs using the 1D, one-hot, and square encoding. The AUC values are obtained from an average of 5-fold cross-validation using the mouse datasets.

On the human datasets (i.e. Table VII), the gkm-SVM performed the best in terms of average AUC values for 6 out of 10 of the datasets. However, Basset obtained best average of (0.91) of all the human datasets. Generally, the results are quite mixed for the human datasets since the best predictors for different datasets are distributed to all the methods used. However, clearly, the one-hot with CNN does not perform as good as other methods using the the mouse datasets. Fig. 3 shows the average AUC values for all the compared methods.

**Table VI.** Comparison of the average aucs using different sequence encoding methods with cnn versus basset and gkm-SVM on the mouse datasets

| Mouse | 1D | 3Layer | One-hot | Square | Basset | gkm-SVM<sup>*</sup> |  |
| --- | --- | --- | --- | --- | --- | --- | --- |
|  |  |  |  |  |  | Complement | NullSeq |
| BCL5 | 0.9980 | **0.9990** | 0.9988 | 0.9985 | **0.9991** | 0.7602 | 0.6695 |
| CBP | 0.9406 | 0.9351 | **0.9473** | 0.9284 | **0.9496** | 0.8824 | 0.6621 |
| CDX2 | **1.0000** | 0.9989 | **1.0000** | 0.9989 | **1.0000** | 0.9747 | 0.8152 |
| CRX | **0.9651** | 0.9403 | 0.9592 | 0.9429 | 0.9630 | 0.8377 | 0.8022 |
| ERG | 0.8834 | 0.7883 | 0.8546 | 0.8635 | **0.8913** | 0.8062 | 0.8892 |
| ETS1 | 0.9408 | 0.9343 | 0.9126 | **0.9461** | **0.9542** | 0.9308 | 0.9332 |
| EZH2 | **1.0000** | **1.0000** | **1.0000** | **1.0000** | **1.0000** | 0.8354 | 0.6478 |
| FLI1 | 0.8442 | 0.7430 | 0.8637 | 0.8168 | **0.8974** | 0.7923 | 0.8922 |
| GATA2 | 0.8932 | 0.7826 | 0.8603 | 0.8755 | **0.9342** | 0.8670 | 0.8924 |
| GFI1B | 0.8623 | 0.7866 | 0.7986 | 0.8323 | **0.9004** | 0.8387 | 0.8774 |
| Average | 0.9328 | 0.8908 | 0.9195 | 0.9203 | **0.9489** | 0.8525 | 0.8081 |

\* The “complement” uses the complement datasets as the negative datasets; while “NullSeq” uses gkm-SVM null sequence function to generate the negative sequences.

**Table VII.**
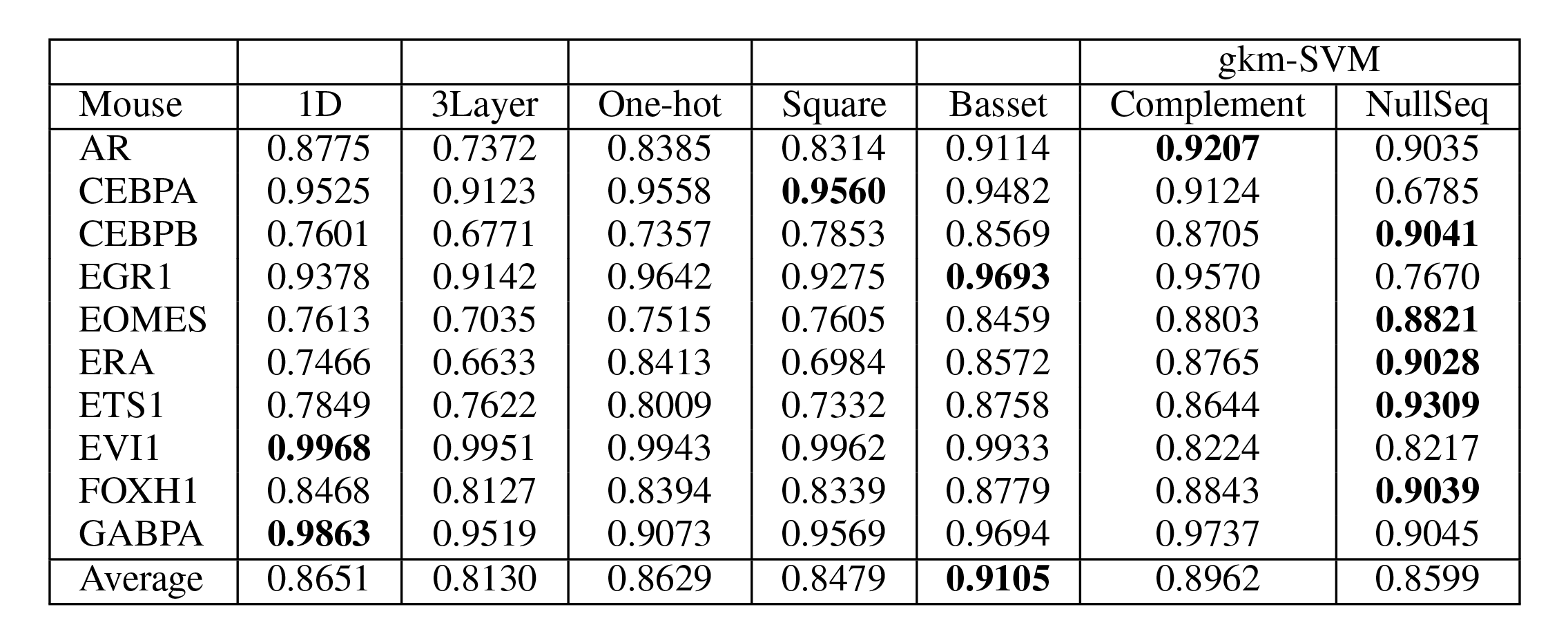
Comparison of the average aucs using different sequence encoding methods with cnn versus basset and gkm-SVM on human datasets

**Fig. 3.**
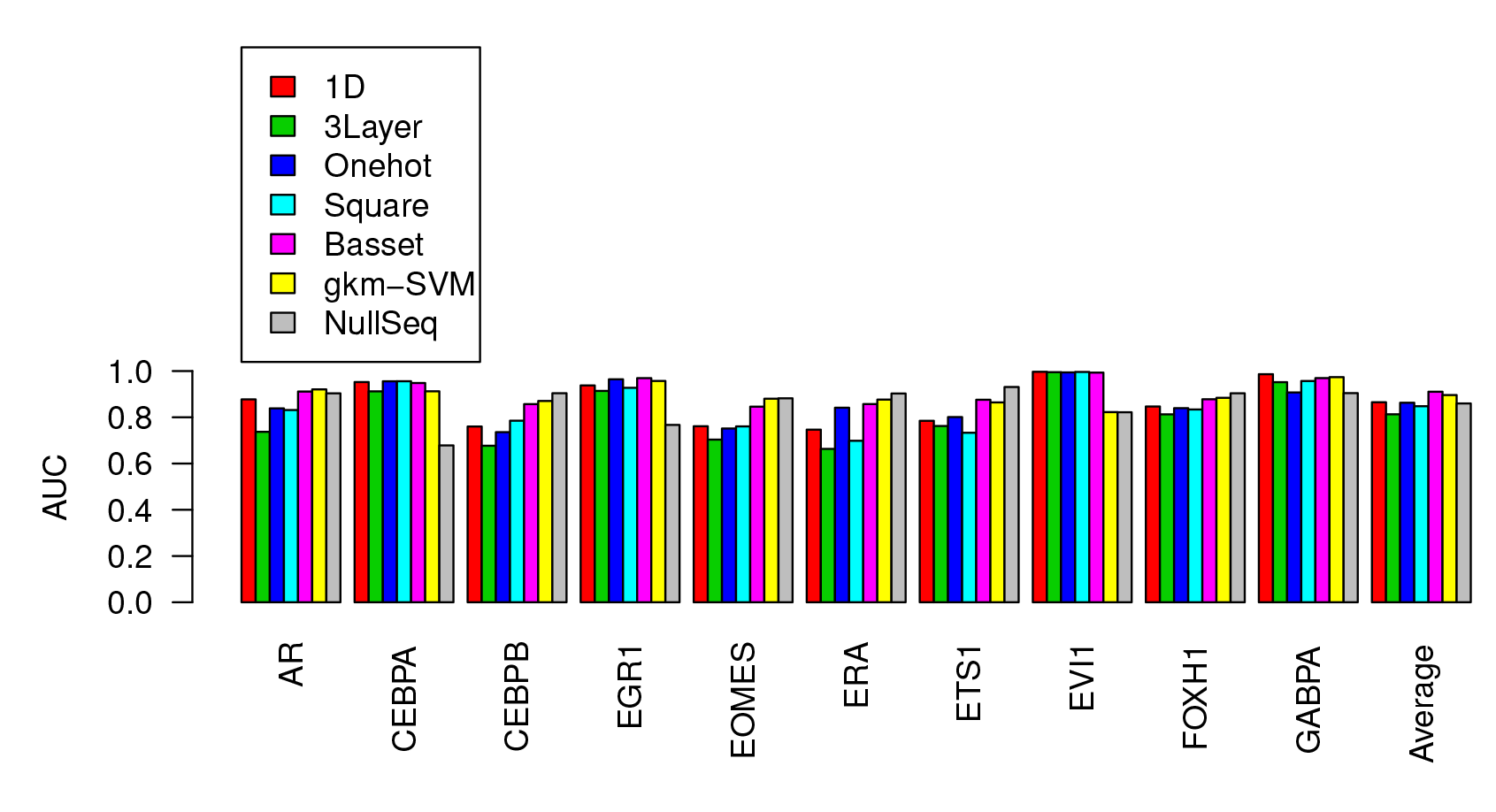
Comparison of the CNN average AUCs using the 1D, one-hot, and square encoding. The AUC values are obtained from an average of 5-fold cross-validation using the human datasets.

## IV. Discussion and Conclusion

This study investigated how a simple ordinal encoding method would perform in comparison with the state-of-the-art one-hot encoding method for DNA sequences. Specifically, the encoded sequences are meant for CNN learning which expect the input examples are in vector or matrix format. What are the desired properties of an encoding method for CNN to model effectively the DNA sequences enriched with motifs? That question is challenging because it would require domain experts with the understanding of the relevant features for the prediction of various classes of motifs. For instance, it was discovered that the frequencies of the short sequences (k-mers) in ChIP sequences are useful for the construction of classification model [19], [20]; while for enhancers it was found that the short repeats are important for their location identification [21]. In addition, for predicting cis-regulatory modules (CRM), the existent of a set of motif signals clustered within a pre-defined region length in DNA sequences are useful signature for their identification [21]. Other than that, for histone marks which are associated with active enhancers, previous studies found no clear sequence patterns that can be signals for their identification [22]. However, past studies have demonstrated that the k-mers are discriminating features for various histone marks [23], [24] and ChIP sequences [19]. The one-hot encoding method which assumes the features in DNA sequences can be detected by filters (i.e. PFMs) in the convolutionary layers may not be able to detect those features mentioned. Therefore, one-hot encoding may not detect distinctive features in different means of DNA sequences.

Our method is termed direct encoding as opposed to indirect encoding [9]. The ordinal encoding is a direct encoding method which preserved the original order and position of each nucleotide in a DNA sequence. Even after the 1D vector is reshaped, the nucleotides ordering is still preserved in the resulted matrix but in a different locations (e.g. rows). The CNN would still be able to detect those features because of the filters scanning.

Our evaluation results using various datasets showed that the ordinal encoding with square matrix representation has good performance on ChIP datasets. Nevertheless, Basset, which is using the one-hot encoding has optimized architecture to achieve good performances across all the datasets. While that is the case, the results of the ordinal encoding with matrix representation are quite close to Basset (one-hot) for the Mouse and Human datasets. The advantage of the ordinal encoding is its shorter input dimension which reduces the computations of filters scanning in the convolutionary layer. Likewise, the kernels in the pooling layer would require less computations.

Table VIII shows the total running time for the three sets of datasets for training the CNN classifiers. The simulation ran on a Linux PC, with 8GB RAM, iCore7 (2.5GHz) processor and 4GB NVidia GeForce 930M graphic card. It is clearly seen that the Square encoding has saving of almost one third of the running time in comparison to one-hot and 1D encoding method. The 1D and one-hot encoding methods have almost the same running time because the number of computations required at both pooling and convolutionary layers grows with the length of matrix (vector). Because the implementation is different in terms of programming language and data structure between Basset (using Torch7 and Lua) and TensorFlow (Python), it is hard to compare the running time for both tools. Basset took 17m20s for the ChIP-seq datasets, while are 29m11s and 29m2s for the Human and Mouse datasets, respectively.

**Table VIII.**
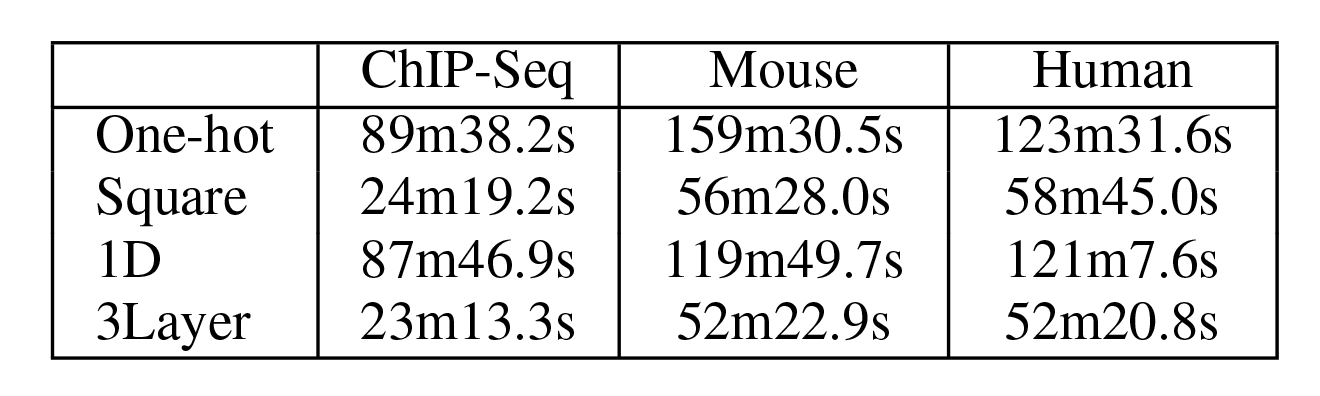
Total CNN running time used for chip-seq, mouse, and human datasets using cnn implemented by tensorflow.

|  | ChIP-Seq | Mouse | Human |
| --- | --- | --- | --- |
| One-hot | 89m38.2s | 159m30.5s | 123m31.6s |
| Square | 24m19.2s | 56m28.0s | 58m45.0s |
| 1D | 87m46.9s | 119m49.7s | 121m7.6s |
| 3Layer | 23m13.3s | 52m22.9s | 52m20.8s |

Our evaluation results also demonstrated that the 3Layer CNN does not perform well for the square representation. According to [25], CNN architecture decision is task-specific and therefore, further tuning may necessary to find the suitable architecture configuration.

The main conclusions from this study are:

- The ordinal encoding method for DNA sequences performs comparably to the one-hot encoding. While the one-hot encoding has better interpretation, it has not much computational and performance advantages over the simple ordinal encoding. This study did not investigate the convergence behavior of both encoding methods.
- The reduced input dimensions using the ordinal encoding allows CNN to learn the input examples with reduced time. However, the speedup gained is depends on the implementation of the CNN tool.
- Different encoding and representation methods may require customize tuning of CNN architecture and parameters used. Currently, there is still lacked of guidelines on choosing those parameters for learning DNA sequences.

